# Lateral phase differences in a population model of the visual cortex are sufficient for the development of rhythmic spatial sampling

**DOI:** 10.1101/2020.07.11.198820

**Authors:** Justin D. Yi, Katsushi Arisaka

## Abstract

When attending to many spatially distributed visual stimuli, attention is reweighted rhythmically at 4-8 Hz. The probability of detection depends on the phase at which a stimulus is deployed relative to this intrinsic rhythm. The reweighting oscillations can be observed both electrophysiologically and behaviorally, and appear to be regulated by the pulvinar. Based on these findings, we considered the computational consequences of allowing feedback to shape the distribution of inhibitory oscillations from the thalamus, as measured by a local field potential (LFP) phases in the 8 Hz low alpha-band, across laterally-connected regions of the visual cortex. We constructed a population activity model with lateral and feedforward connections. In agreement with prior models, we found that the sign of the lateral phase difference in the inhibitory low-frequency oscillations regulated the direction of communication between the laterally-connected regions. Furthermore, the phase difference induced periodicity in the dynamics of a downstream winner-takes-all attractor network such that periodic switching between states was observed. We finally simulated a simple spatial attention task. We found rhythmic 8 Hz sampling between two regions when a lateral phase difference was present—an effect that disappeared when the lateral phase difference was zero. These findings are in agreement with spatial attention literature and suggest that lateral phase differences are essential for manifesting communicational asymmetries in laterally-connected visual cortices. Our model predicts that population-specific phase differences are critical for sampling the spatial extent of stimuli.

**Author summary:** We conducted a computational study of the effects of lateral phase differences in a simulated model of the visual cortex. Lateral phase differences are defined to be when the phase of an intrinsic low-frequency inhibitory oscillation varies consistently across populations in the same cortical area. For example, our model was intended to capture the dynamics of a retinotopic cortex where feedback from the frotoparietal areas via the pulvinar nucleus assigned laterally-connected regions of the visual cortex different phases. We found that the sign of the phase differences influenced the direction of lateral communication. Furthermore, the phase differences introduced rhythmicity in the downstream areas, thus allowing us to simulate rhythmic spatial selection of stimuli. Prior to the current study, the influence of inter-areal phase differences in feedforward models had been well characterized. Our model provides new insights into the dynamics of population-specific lateral phase differences and predicts that the development of phase differences across the visual cortex are critical for the allocation of attention in space.

## Introduction

Neural oscillations in the visual system are capable of facilitating a vast array of computational functions. Models of visual computation must take into account oscillatory interactions across space and among a broad distribution of frequencies. The theory of Communication Through Coherence (CTC) [1] was developed to explain how temporal windows of interregional communication develop in the presence of oscillations. CTC predicts rapid synchronization in the bottom-up gamma band is regulated by top-down alpha frequency rhythms [2]. Furthermore, neural oscillations have been implicated in rhythmic spatial attention via low-frequency theta and alpha oscillations coordinated by the frontoparietal regions [3–6]. This growing body of literature suggests rhythmicity is essential for the allocation of resources to, and the processing of, relevant visual signals.

The interactions between low- and high-frequency bands shape communication in the visual system. The main effector of low-frequency band feedback seems to be the broadly-projecting pulvinar nucleus (PN) of the thalamus [7, 8], whereas gamma-band oscillations may emerge from the coupled excitatory-inhibitory interactions between pyramidal cells and interneurons in the cortex (for a review, see [9]). The phase of low-frequency cortical oscillations, potentially organized by the thalamus [10], regulate windows of communication, excitability, and directionality of transmission in the high-frequency bands [11–15].

Provocative questions emerge when considering the evidence for retinotopic distribution of low alpha-band activity in the visual cortex. It has been demonstrated with magnetoencephalogram recordings during a spatial attention task that task-relevant changes in alpha power correspond to the cortical retinotopic map [16]. Furthermore, stimulus cross-correlation analysis against occipital electroencephalogram traces has shown that stimulus-induced 10 Hz perceptual echoes emerge across the cortex as traveling-waves [17]. Given this evidence, it is clear that the spatial extent of the phase of low alpha-band oscillations is of critical importance for neural encoding schemes of visual space [18].

What are the computational consequences for a neural system with low-frequency phases differentially distributed over the retinotopic map? What is the effect on the transmission through lateral cortical projections, and between subsequent downstream cortical areas with analogous representations of retinotopic space? We hypothesize that the spatial distribution of phases of low-frequency rhythms are essential to process the spatial extent of stimuli.

Previous models of encoding have made similar predictions that the phase of intrinsic rhythms is critical to encode space. For example, an interpretation of hippocampal theta phase precession in rodents proposes that phase coding facilitates a neural representation of the world by a sequence of events without invoking space or time [19]. Another model of encoding in the visual system predicts that, in principle, a visual stimulus can be encoded and decoded reliably by neuron-specific phase shifts in high-frequency oscillations [20]. These studies support the notion that population-specific phases are utilized by the brain to represent space. We seek to extend this body of understanding by elucidating how assigning spatial specificity to lateral phase differences in the low alpha-band shape the dynamics of spatial attention.

We choose to investigate how population-specific retinotopic distributions of the low alpha phase temporally segment gamma-band transmission and promote temporal scansion [21] between stimuli. In this proposed model system, functional communicational asymmetries in lateral communication within the same cortical area modulate the selectivity of downstream cortical regions via temporal multiplexing. We assume that the decision to attend to or process a stimulus is governed by a winner-takes-all attractor [22–25], and that competition through mutual inhibition allows for the rhythmic processing of multiple stimuli [26]. We also investigate how the rhythmic selection of stimuli based on the phase difference across lateral connections in our attractor model may extend the findings in the literature on the nature of rhythmic spatial attention.

## Materials and methods

### Model architecture and dynamics

To investigate the excitatory-inhibitory dynamics of the visual cortex, we used a non-linear mean field population activity model inspired by the Wilson-Cowan equations [27, 28].

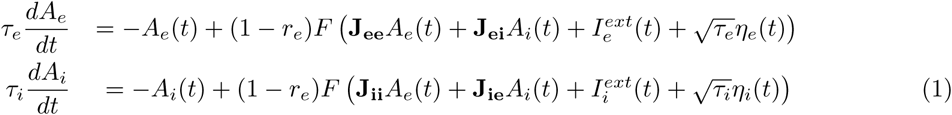

*A*_*n*_ describes the dimensionless activity, or mean firing rate, of a population of *n*-type neurons where *n* ∈ [*e, i*]. The subscript *e, i* denotes whether the population is excitatory or inhibitory. The coefficients *r*_*e,i*_ describe the proportion of neurons in the absolute refractory period for a given time step, and were given relatively small values of 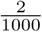 and 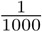, respectively. The matrices **J_nm_** describe the connection strengths between population *m* projecting to population *n. A*_*e*_ and *A*_*i*_ were further decomposed into vectors of length 4 and 3, respectively, to encode the activity (and their derivatives) of subpopulations E1-4 and I1-3 (Fig. 1).

**Fig 1.**
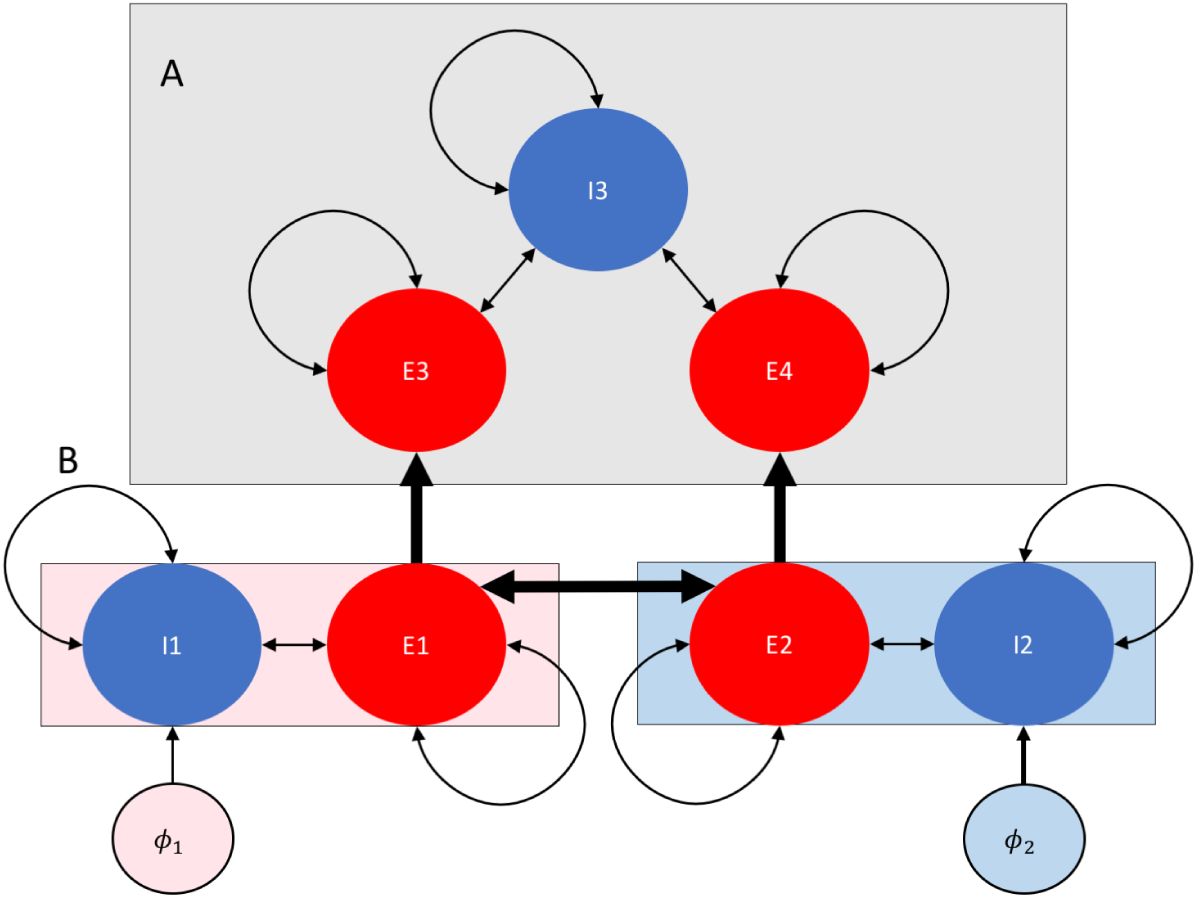
Schematic of model network with lateral phase gradient. Excitatory subpopulations E1-4 and inhibitory subpopulations I1-3 are colored red and blue respectively, with connections represented by arrows. **(A)** E3 and E4 share inhibition with I3, which is responsible for establishing competition between the excitatory subpopulations leading to attractor dynamics [22]. **(B)** E1 and E2 project in a feedforward manner to E3 and E4. E1 and E2 share reciprocal lateral connections where the activity in I1 and I2 is modulated by sinusoidal oscillations with phases *ϕ*_1,2_.

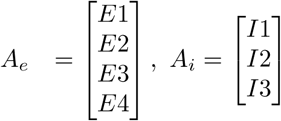

To couple the subpopulations described by these activity vectors and their derivatives, matrices were used to represent the sign and strength of connections between and within the subpopulations. They were defined as follows:

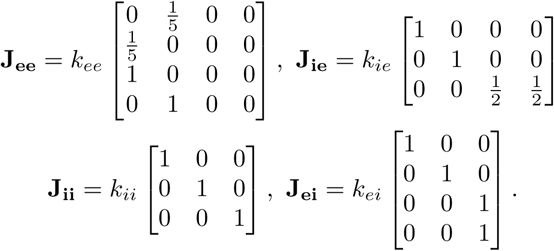

The scalar coefficients in front of each matrix took values of *k*_*ee*_ = 1.5, *k*_*ie*_ = 3.5, *k*_*ii*_ = −2.5,*k*_*ei*_ = 3.25 a.u. (arbitrary unit amplitude) in order to replicate the frequency spectrum of activity in layers 2/3 of the cortex [12].

*τ*_*n*_ describes the membrane time constants of the population *n* (6 ms for *n* = *e* and 15 ms for *n* = *i*), 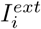 describes the external driving current, and *η*_*n*_ describes the Gaussian white noise that the population received with variance 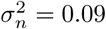 = 0.09. We define *F* (*x*) to be a rectified linear activiation function:

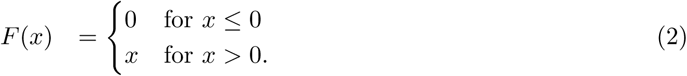

A schematic diagram of the circuit diagram can be seen in Fig. 1. An attractor network was constructed from two excitatory populations competing through shared inhibition (Fig. 1A). We assume this attractor is representative of higher order visual areas responsible for the selection of relevant stimuli [25]. We feed this attractor with input from a highly simplified retinotopic cortex, representative of cortical columns on V1, in which only two populations exist, each selective to a different stimulus location (Fig. 1B). In this low-order region, only Regions 1 and 2, each with subpopulation of excitatory (E) and inhibitory (I) cells, received a direct-current injection of either 0.5 (“low” condition) or 1.5 (“high” condition) a.u. to the E cells, and an alternating-current (AC) to the I cells to simulate a coherent 8 Hz inhibitory input from the pulvinar (Fig. 1B)[15]. We assume that the phase of synchronized inhibitory activity from the pulvinar can be measured by the local field potential in the region in question and is also reflected in the cortical activity modulation. Of note is that feedforward, recurrent, and lateral connections were all considered in this model. The dynamics of the system were studied for when the phase of the AC currents was different across the lateral regions, i.e. when the relative phases shifts were of 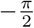, or 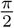 radians.

### Simulated inhibitory oscillations

We investigated the model responses when the AC input to subpopulation *n*, 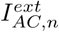, was the sum of sine components of different frequencies with the respective magnitudes *M*_*AC,i*_. The AC components were used to drive the inhibitory populations with a low-frequency alpha rhythm assumed to originate from the pulvinar nucleus (PN) of the thalamus [13, 15] to simulate an oscillation of coherent inhibitory activity with a mean frequency of ∼ 8 Hz. To capture a realistic qualitative amplitude distribution of frequency components, the oscillations were constructed from the sum of sine waves where the amplitudes were governed by a 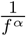 distribution summed with a Gaussian centered on 8 Hz.

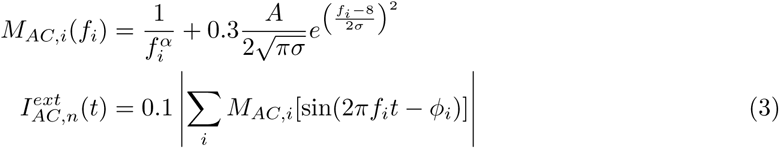

The value *A* was set to have a value of 10, *σ* was 0.5 Hz, and the range of frequency components *f*_*i*_ considered was 0.1 − 200 Hz. The parameter *α* was chosen to be 0.4 to give more weight to the high frequencies to serve as noise (Fig. S1). The phase of each frequency component, *ϕ*_*i*_ was drawn from a uniform distribution of [−*π, π*]. In order to calculate a phase shift of Δ*θ* with some jitter, the original phase components *ϕ*_*i*_ were shifted by the rule *ϕ*_*i*_ − (0.3ζ + Δ*θ*). ζ was a random number drawn from a uniform distribution of [− *π, π*] radians, then rescaled by a factor of 0.3 to introduce jitter. However, if a phase shift of exactly Δ*θ* = 0 was to be considered, the jitter coefficient was set to 0.

Before the oscillations were injected into an inhibitory population, the amplitude was rescaled to have a maximum value of 1 a.u., indicated by the absolute value bars in Eq. 3, then multiplied by an amplitude value of 0.1 a.u. (however, for Fig. 2 and 3C, it was 1.0 a.u.). All simulations were run with a time step of Δ*t* = 0.1 ms and numerically integrated with Euler’s method (except during Coherence and Granger Causality analysis, where Δ*t* = 1.0 ms).

**Fig 2.**
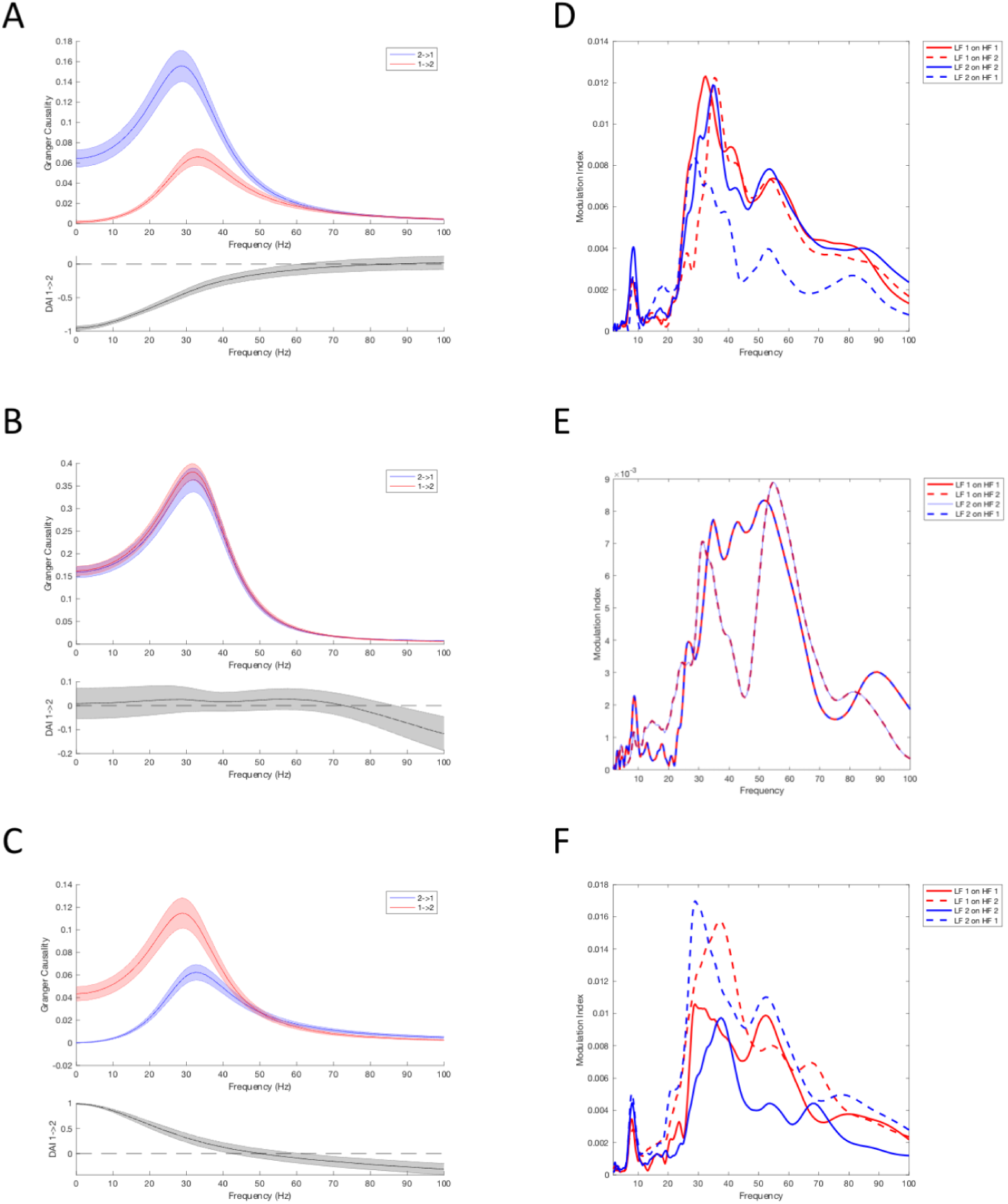
Lateral phase gradients induce asymmetric lateral communication. After 30 trials were conducted, the Granger Causality and Directed Asymmetry Indices **(A-C)**, and the modulation index (MI) for phase-amplitude coupling (PAC) **(D-F)** were calculated. The top row **(A**,**D)** was for the condition where 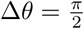 (see Materials and Methods), the middle row **(B**,**E)** was for Δ*θ* = 0, and the bottom row **(C**,**F)** was for 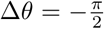 radians. It is evident that the sign of lateral phase difference inverted the directionality of lateral communication. The phase of low-frequency oscillation modulated the envelope of the 20-60 Hz frequency components most strongly **(D-F)**. In **(A-C)**, the shaded error bars represent the standard deviation calculated from the bootstrapped PWCGC.

**Fig 3.**
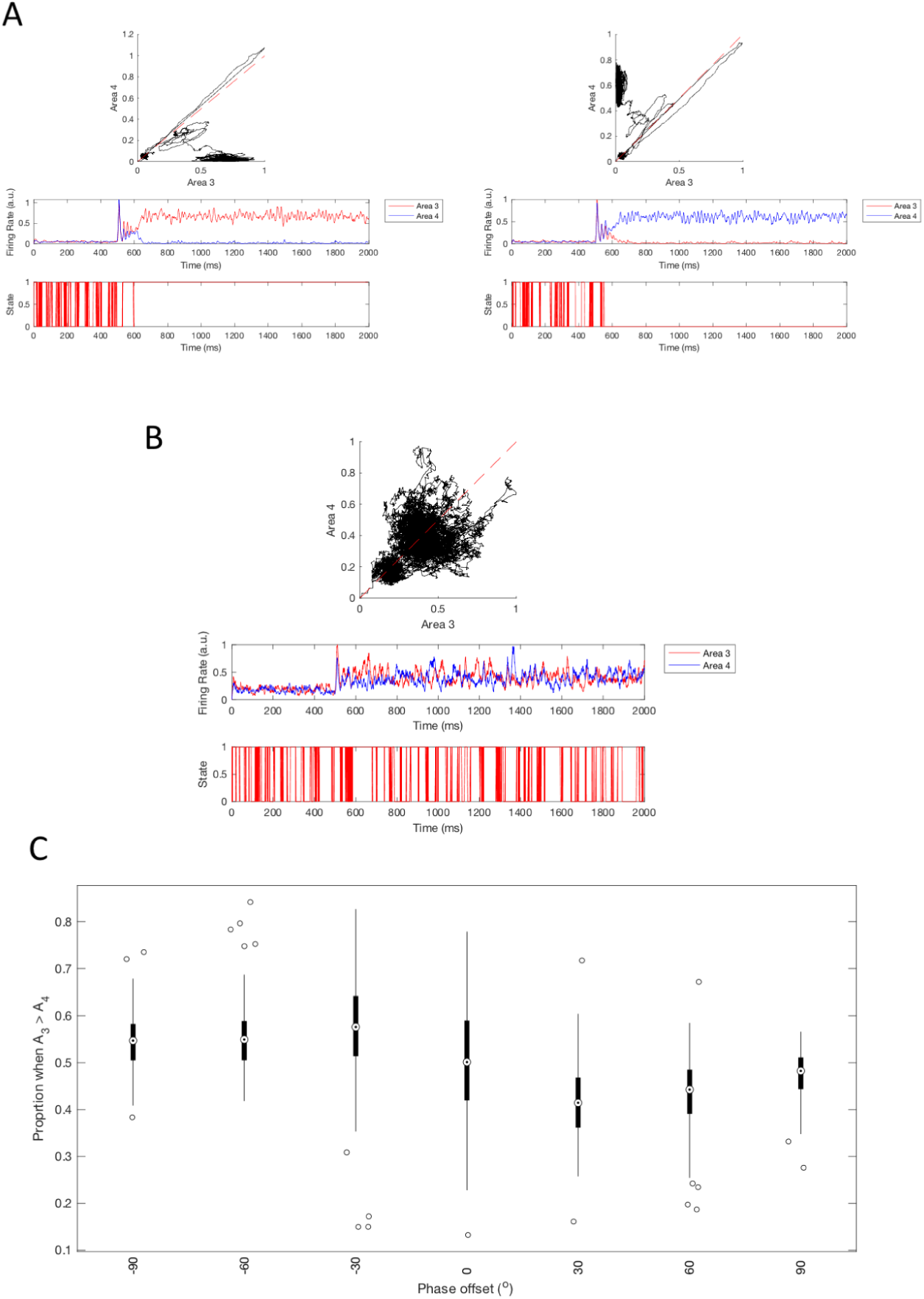
Attractor dynamics are modulated by lateral phase differences. **(A)** When the stimulus supplied to E1 and E2 was equal and high (turned to 1.5 a.u. at 300 ms), the attractor dynamics were probabilistic and exhibited winner-takes-all behavior [22], even when E1 and E2 had a phase difference of 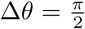. The phase planes show the two cases where E3 (left) or E4 (right) “wins.” The dashed red line indicates *x* = *y*. Beneath the phase planes are the time series of the activity of E3 and E4. The “State” refers to when the activity in E3 > E4. **(B)** When the stimuli were low (turned to 0.5 a.u. at 300 ms), then there was weakly periodic switching of the “State” as a function of time. **(C)** 100 trials were conducted for each phase difference and the probability that the “State”= 1 was found from *t* > 300 ms for each trial. The magnitude of the phase difference systematically shifted the probability that the State was E3 > E4 as a sinusoid when the oscillation was allowed to take a larger value (1 a.u., compared to the usual 0.1 a.u. defined in the Materials and Methods). The box-and-whisker plots show the median (dot), Q1 and Q3 (boundaries of thick blocks), and maximum and minimum (thin tails), excluding outliers (hollow dots).

### Coherence analysis

The multi-taper coherence between the *A*_*e*_ continuous time series for Regions 1 and 2 (E1 and E2) was calculated using the CHRONUX toolbox for MATLAB [29,30]. From 30 trials where the oscillations were re-drawn for each trial, jackknife error bars were calculated, assuming a significance level of *α* = 0.01. The fast Fourier transform (FFT) was padded with the setting of “1”. *K* = 5 tapers used were defined to have a time-bandwidth product of *TW* = 3. The sampling frequency was set to be 1 kHz based on the time step, Δ*t* = 1.0 ms, used for numerical integration of the simulation.

### Granger Causality (GC) and Directed Asymmetry Index (DAI)

The pairwise conditioned Granger causality (PWCGC) between the *A*_*e*_ continuous time series for Regions 1 and 2 (E1 and E2) was calculated using the Multivariate Granger Causality toolbox for MATLAB [31]. The PWCGC was calculated by a bootstrap method, sampled from 30 trials where the oscillations were re-drawn for each trial, to find the conditioned spectral Granger causality (GC) from an estimated auto-covariance matrix. The number of lags used was set to be 5, approximately the value found by the Bayesian Information Criterion model estimation. The PWCGC function returned the GC interaction strengths for both the E1 → E2 and E2 → E1 directions. To reduce computation times, the numerical integration was performed with Δ*t* = 1 ms, and to match the unimodal shape of the GC distributions in prior literature [12, 15], the amplitude of the inhibitory oscillation was increased to 1.0 a.u. from the usual 0.1 a.u.

From the GC in the two different directions for a given frequency component, the Directed Asymmetry Index (DAI) [12, 32] was calculated along the E1→E2 direction:

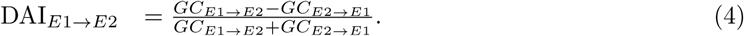

The mean and standard deviation of the GC and DAI for each frequency component across the 30 trials were calculated.

### Phase-Amplitude Coupling (PAC)

The phase-amplitude coupling (PAC) for the instantaneous phase of the low-frequency oscillation, *ϕ*_*LF*_ (*t*), and the envelope, *a*_*HF,i*_(*t*), of a given high-frequency component, *f*_*i*_, was quantified using a modulation index (MI) [33, 34]:

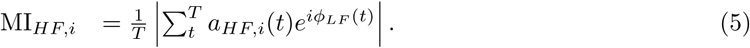

*ϕ*_*LF*_ (*t*) was extracted from the instantaneous phase of the analytical signal of the low-frequency oscillation found by the Hilbert transform. The envelopes of the *A*_*e*_ signals, *a*_*HF,i*_(*t*), were recovered from the absolute value of the narrowband signal with frequency *f*_*i*_, found by the continuous wavelet transform (CWT) in MATLAB. The CWT used 48 voices per octave. The PAC was evaluated for low-frequency oscillations and HFs within and across Regions 1 and 2.

### Simulated Spatial Attention Task

A spatial attention task [4] was given to the model network. To simulate this task, the two Regions 1 and 2 were assumed to encode different locations in space. A relative phase difference of 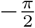, 0, or 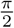 radians was assigned to Regions 1 and 2. Only for when the network had a 0 radian phase difference, we set the phase shift jitter value ζ = 0 in order for the phase difference to exactly be zero. We considered the network to be in the “correct” state to “see” a stimulus at time *t* when the instantaneous activity was E3 > E4 for a given t=300-1100 ms. This definition of only considering E3 > E4 is valid because the network could not “predict” where the stimulus would appear based on prior trials, so presenting the stimulus repeated at the same “spatial” location (i.e. testing for only E3 > E4) was equivalent to if the stimulus position was varied randomly (as would be the case in a real psychophysical test).

The network was stimulated at *t* > 300 ms with a value of low 0.5 a.u. (high 1.5 a.u. was tested in Fig. S2). This stimulation is not to be thought of as the stimulus presentation, but rather the network being “cued” or “primed” to rhythmically search the spatial location via covert attention. The time to sample the state of the network was selected in 20 ms intervals, and 30 repetitions were conducted for each sampling time. The simulated inhibitory oscillations were kept constant across sets of 30 repetitions and only changed when the different phase differences were tested. To quantify the rhythmicity of target selection, the FFT of the detrended time series of the proportion of “correct” trials was calculated.

To test whether or not the amplitude of an observed rhythmic fluctuation in the proportion of “correct” trials was significant, a bootstrap statistical test of significance was conducted [4]. 1,000 random permutations of the proportion of “correct” v.s. “incorrect” trials across all 30 trials and all times tested were configured and converted into 1,000 detrended time series of proportions. From this bootstrapped sampling distribution of time series, the FFTs were calculated and for each frequency bin, the 95-percentile amplitude was figured and used as the boundary for significant amplitude.

All analysis and numerical integration of the activity models was performed using MATLAB Version 2019b [35]. Shaded error bars were constructed with the shadedErrorBar function [36].

## Results

### Lateral phase differences create communicational asymmetries

We studied our model in the case where there was a phase difference in the simulated 8 Hz inhibitory oscillations between laterally-connected Regions 1 and 2 (Fig. 1). In the cases where the phase difference between the regions was 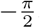, 0, or 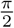 radians, we noted that the coherence between the regions was most prominent for the ∼ 8 Hz components and the 15-35 Hz components (Fig. S2). Furthermore, the sign of the phase difference dictated the direction of lateral communication as measured by the PWCGC and DAI (Eq. 4). For a positive phase difference, Region 2 led Region 1 in frequency components <50 Hz (Fig. 2A). For a negative phase difference, the opposite directionality was observed (Fig. 2C). For zero phase difference, neither Region led the other Fig. 2B).

Our finding that the phase of the underlying low-frequency oscillation determined the direction of communication is consistent with other models of the interactions between different areas of the visual system [12–15]. We also observed that the phase of the inhibitory oscillation was preferentially coupled (via the MI from Eq. 5) to the amplitude of frequency components in the 20-60 Hz range (Fig. 2E,F). The phase of oscillations were coupled to the high-frequency components both within the same Region and across different Regions 1 and 2. We note that when the phase difference was exactly zero, the high frequencies of both Regions were equally coupled to the phases of the low-frequecy oscillations originating from either Region (Fig. 2E). PAC is a phenomenon observed in many models of interacting neural oscillations [12–15,32–34,37]. However, our model predicts that asymmetric communication and PAC is expected to be observed between laterally-connected regions in the same visual area if the phases of the underlying inhibitory oscillations are population-specific.

### Lateral phase differences promote rhythmicity in downstream attractors

In our model, the Regions 3 and 4 shared inhibition and thus competed. If Regions 3 and 4 were driven by strong input to Regions 1 and 2 (Fig. 3A), the mean firing rates showed well-known winner-takes-all attractor dynamics where only one region was allowed to fire [22–24]. We define a “State” to quantify when the activity of excitatory Region A3 exceeded that of excitatory Region A4 (notated as E3 > E4).

In the high-intensity input regime (1.5 a.u.), the winner-takes-all dynamics superseded the intrinsic rhythmicity of the activity of Regions 1 and 2 (Fig. 3A). However, if the input was of low-intensity (0.5 a.u.), then the dynamics were no longer winner-takes-all and spontaneous switching between States was weakly periodic (Fig. 3B). In fact, the magnitude of the phase difference between Regions 1 and 2 dictated the proportion of the time that the network was in State = 1 (Fig. 3C). Over 100 trials for each phase difference, the median proportion of time points that State = 1 was observed to varied as a sinusoid. This finding demonstrates that lateral phase differences can disrupt the attractor dynamics and create a functional asymmetry where one State is sampled, on average, more frequently than the other.

We argue the State in which the attractor network is in can be interpreted as whether the system is susceptible to input at a given location in space, i.e. whether that location is sampled by attention at a given time point [3, 4, 6, 25]. Therefore, we simulated an analog of a simple spatial attention task using our model network, assuming the 0.5 a.u. input drove intrinsic 8 Hz rhythmicity in the State, governed by the phases of the oscillations (Fig. 4A). For our simple example, we probed only one “spatial” location of the network by asking if State the network was E3 > E4 for a given time. We argue that sampling the state of the network is analogous to presenting a stimulus at that time and probing whether or a subject detected it.

**Fig 4.**
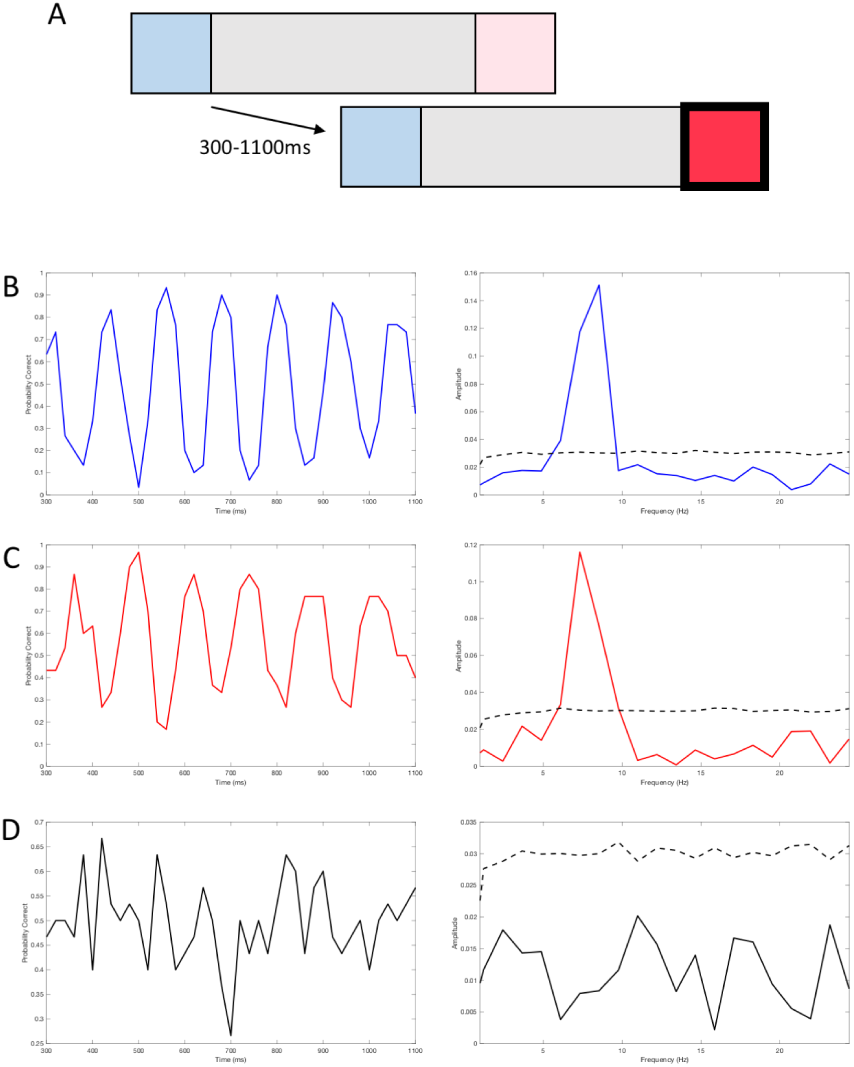
Rhythmicity in attractor dynamics recapitulates rhythmic spatial attention. **(A)** A schematic of a simple spatial attention task where a subject must indicate where a change in the contrast occurs after a variable delay (compare blue versus red). We assume that there is a spatial extent to the lateral regions of the cortex monitoring the blue or red region. To simulate an analogy to this attentional experiment, we counted the probability that the network was in a “State” where E3 > E4 while the network rhythmicity was activated an input of 0.5 a.u. for *t* > 300 ms. This input can be thought of as the network being activated to “search” between the two states under the direction of attention. **(B)** The left plot shows the probability after 30 trials, sampled for t=300-1100 ms in intervals of 20 ms, that the State was = 1 (i.e. E3 > E4). The right plot shows the fast Fourier transform (FFT) of the selection signal. The dashed line indicates the bootstrap estimated significant amplitude (see Materials and Methods). For a 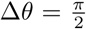, the dynamics of the selection process showed heightened selectivity with a frequency of ∼ 8 Hz. **(C)** When 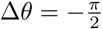, the rhythmic selection was also significant **(B)**. This finding qualitatively replicates the psychophysical results of rhythmic spatial attention, where sampling between the cued and un-cued ends of an object was modulated at ∼ 8 Hz [4]. When there was no phase difference between the lateral regions, **(D)** no rhythmic sampling occurred at all. Note that the cross-condition of relative phase differences here does not necessarily govern the relative phase relationship between the sampling rhythms.

In a qualitative sense, the results from our simulated attentional task agree with prior psychophysical findings in the area of rhythmic spatial attention. We found ∼8 Hz rhythmicity in the sampling between the two spatial locations. When the phase difference was 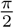 (Fig. 4B) and when it was 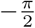 (Fig. 4C), the rhythmicity in the sampling between regions was of significant amplitude. However, comparison across conditions of different phase differences (i.e. 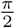 versus 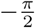 radians) was not predictive of the phase relationships between sampling rhythms since the phase shift value described a systematic shift on the oscillation constructed from random phase components.

Our model’s performance in this simulated attention task was broadly agreement with experiment. A similar qualitative result, although of smaller magnitude, was observed in an analogous spatial attention task with human subjects [4]. In extension of these results, we observed that if no lateral phase difference was present (Δ*θ* = 0 radians), then the rhythmic sampling disappeared (Fig. 4D). The present simulated experiment demonstrates how lateral phase differences promote rhythmic reweighing of spatial attention. However, we note that this solution only exists in our model for low-intensity input (0.5 a.u.) to Regions 1 and 2, since high-intensity input (1.5 a.u.) led to the abolishment of rhythmic sampling and return to winner-takes-all attractor dynamics (Fig. S3).

## Discussion

### Summary of results

Our simulations have elucidated several potential functions that lateral cortical phase differences in low-frequency inhibitory oscillations [15] may serve. Firstly, our measurements of coherency, GC, DAI, and PAC indicate that the low-frequency phase gradients across lateral cortical connections determine an axis of communicational asymmetry. Along this axis, certain regions lead the others depending on the sign of the phase difference (Fig. 2 and S2). The phase differences in the low-frequency oscillations of each region determines the directionality of lateral communication. Secondly, the phase differences affected the population dynamics of downstream attractor networks by introducing weak periodicity in an otherwise winner-takes-all network (Fig. 3). Importantly, the magnitude and sign of the phase difference biased the amount of time the network spent in one State (Fig. 3C). This bias allowed the network to qualitatively recapitulate rhythmic sampling in a simulated spatial attention experiment (Fig. 4B,C). However, in order for rhythmic spatial sampling between different spatial stimuli to occur, a lateral phase difference had to be present or else no sampling rhythm was observed (Fig. 4D).

### Relationship to prior models and experiments

Our model’s interpretation of lateral phase differences may be able to explain critical questions in spatial attention research, such as providing a specific mechanism for how corticothalamocortical interactions in the alpha-band regulate activity in the attention network. It is known that both thalamic and cortical alpha are critical for understanding the dynamics of the attention network. When the pulvinar interacts with the higher-order areas of the FEF and LIP, attentional engagement has been associated with increased alpha power [7]. This implies that alpha-activity in the pulvinar is important for coordinating higher-order centers of attention. Cortical alpha has also been shown to be a predictor of attentional performance. For instance, several studies have demonstrated that attentional enhancement is often broadly characterized by alpha-band power decrease, and attentional suppression is characterized by alpha-band power enhancement with respect to the location of an attentional target [38–40]. We predict that these changes in power are related to the degree of phase (de-)synchronization in the alpha/theta band. We hypothesize that a decrease in alpha power means that phases at different retinotopic locations are scattered, promoting differential rhythmic sampling of the spatial extent of the stimulus. An increase in alpha power would imply that sampling the spatial extent of the stimulus is task-irrelevant, and thus feedback influences from the frontoparietal regions [5, 6] are not deployed to adjust the phases across the retinotopic map.

A caveat to consider is that our model is a simplification of the attentional network. First of all, we decided not to model the dynamics of the higher-order frontoparietal network [5, 6, 41] which are responsible for spatial attention. Instead, we assumed that this frontoparietal network simply assigned low alpha phase differences via the PN, which mapped the phases to specific regions across the retinoptic map in a one-way manner. This assumption that the generators of alpha in the PN are retinotopic is supported by experiment [16]. We did not model reciprocal corticothalamocortical interactions with the PN either. For instance, the ventrolateral PN influences the dynamics of V4 and IT cells depending on attentional demands, and is required to maintain an active cortex [42]. In this network, V4 leads the PN in gamma-band activity which implies the visual cortex modulates the dynamics of the PN in an attention-dependent manner [42]. Furthermore, it has been shown that V4 feedback onto V1 is needed to observe lateral interactions responsible for the perceptual grouping of line segments, and those lateral interactions are also necessary to increase the strength of feedback [43]. These reciprocal feedback dynamics were omitted in favor of the capturing solely the elementary lateral cortical dynamics brought about by phase differences in the low alpha-band.

Despite the aforementioned simplifications, our model still provides insight into the essential dynamics of how the lateral phase influences communication across the retinotopic map. However, future models certainly would benefit from expanding the architecture of our model to consider how large-scale reciprocal connections between the visual cortex, PN, and frontoparietal regions collectively influence the dynamics of attention.

### Predictions and implications

We make a strong prediction that spatial sampling arises from lateral phase differences influencing downstream attractor dynamics. An alternative explanation for rhythmic sampling between stimuli is that periodic strong inhibition shared between many different “embedded” object representations in the cortex causes a “resetting” of attractor dynamics, linking shifts in attention to ongoing oscillations [44]. While our simulation alone can neither affirm nor rule out this possibility, we argue that lateral phase differences complement the aforementioned theory of attention. Namely, in addition to the predicted synchronization of attentional shifts to oscillations [44], we propose that these attentional shifts should be accompanied by lateral phase differences assigned to the different areas of the retinotopic map sampling the stimuli.

Our model predicts that asymmetries in the direction of lateral communication across the retinotopic map are mediated by population-specific phase differences. One important asymmetry to consider is the preferential V4 gamma synchronization observed when simultaneously stimulating two sites on V1 which converge on a single V4 site [45]. Our model’s GC results suggest that this preferential synchronization in the gamma-band may controlled by the sign of the low-frequency phase differences between V1 sites. If one V1 site leads the other in the gamma-band, this may induce the common V4 site to more strongly synchronize with the leading site.

In a similar vein to preferential synchronization, other perceptual phenomena may be understood through the lens of lateral communicational asymmetries. For instance, rivalry in the perception of different stimuli presented binocularly can be modeled by reciprocal inhibition between populations of neurons selected to one stimulus or the other [46, 47]. Our model predicts that a phase difference across populations selective for one stimulus or the other in a binocular rivalry may be able to explain the transitions in dominance reported in these experiments. Population-specific phase differences would create asymmetries in the reciprocal inhibition among populations encoding the different stimuli; this asymmetry may serve as the basis for the switching and competition between the percepts. Therefore, from our model, we predict that associational spatial hierarchies depend on the phase of low-frequency rhythms across the retinotopic map and that awareness of the spatial extent of a stimulus will degrade if the lateral phase differences between retintopic areas are perturbed.

## Conclusion

Our heuristic computational model provides new insights into how attentional feedback processes can exert their influence via the phase of low-frequency inhibitory oscillations. While prior literature focused on the phase differences between different areas of the hierarchical visual system [11–15], we considered phase differences within the same area of the visual system. We found that the phase difference between two populations is sufficient to create a functional hierarchy of lateral communication. When the areas experiencing a lateral phase difference projected to downstream attractors, the phase differences induced periodic sampling of the attractor’s phase plane. Our model qualitatively captured the dynamics of rhythmic attention [4]. Without a lateral phase difference, rhythmic spatial sampling disappeared. The results lead us to predict the phase differences across a retinotopic cortex can serve as the basis for the development of rhythmic sampling.

## Supporting information

**S1 Fig.**
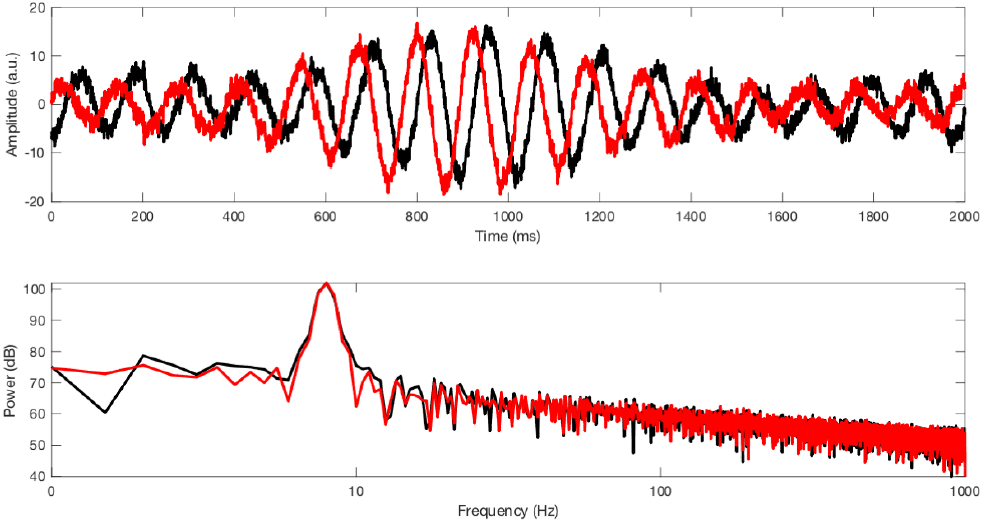
Example of simulated 8 Hz oscillation with 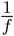 noise profile. Simulated oscillatory signals were constructed from the sum of a Gaussian and 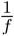 distribution. The top panel shows the signals in the time domain, whereas the bottom panel is the power spectrum of each signal. The relative phase difference between the two signals was 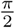 radians.

**S2 Fig.**
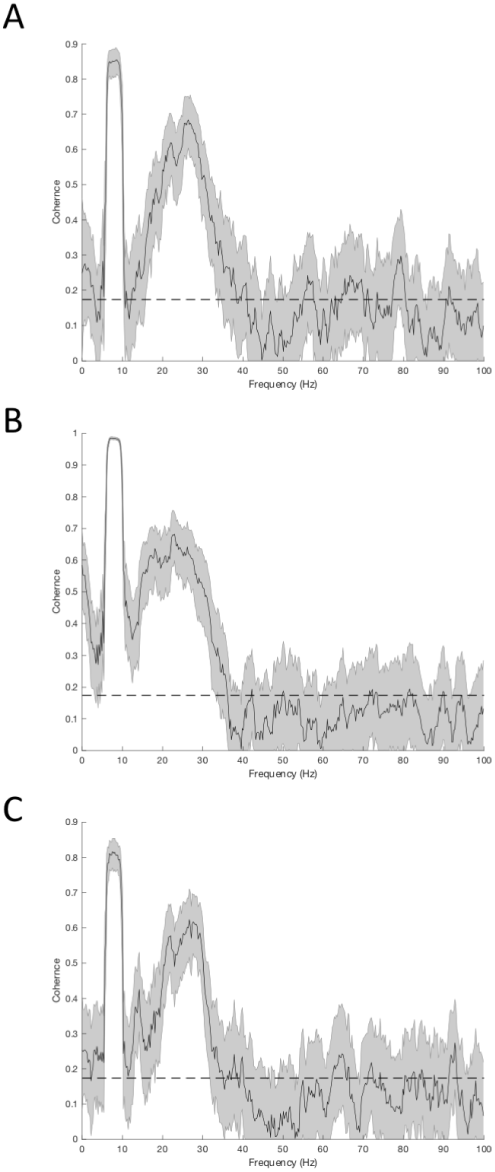
Lateral communication coherence spectra. When Regions 1 and 2 were allowed to communicate with phase differences of 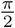 (**A**), 0 (**B**), or 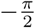 (**C**) radians, the coherence spectra showed prominent peaks at ∼ 8 Hz and 15-35 Hz. Jackknife error bars are shown, as well as a dashed line indicating the *α* = 0.01 significance level.

**S3 Fig.**
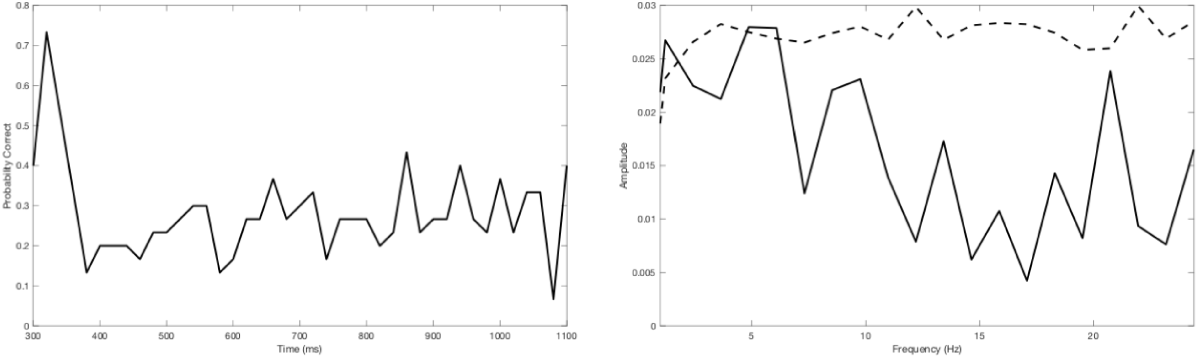
8 Hz rhythmic selection disappears in the high-stimulus limit. A simulated spatial attention task, as in Fig. 4, was conducted but with the priming input to induce rhythmic sampling being “high” (1.5 a.u.). Rhythmicity disappeared because the dynamics became like a winner-takes-all attractor [22], outweighing the effect of the lateral phase differences across the network.

### S1 File. Minimal Data Set and MATLAB Simulation Code

The data to reproduce all findings and figures (provided as .csv files with README documents), as well as the MATLAB code for all the simulations (provided as .m files) are at the following link: https://github.com/jd-yi/Lateral-Phase-Difference-Simulations.

## Acknowledgments

We are grateful to Javier Carmona, Elizabeth Mills, Ikaasa Suri, Saba Doustmohammadi, and Stan Schein for their helpful comments and insightful discussions during the development of this model. We are indebted to Aaron Blaisdell and Zahra M. Aghajan for their expertise and guidance when conceiving of the theoretical framework presented in this work.

